# Trait evolution in adaptive radiations: modelling and measuring interspecific competition on phylogenies

**DOI:** 10.1101/033647

**Keywords:** adaptive radiation, character displacement, phylogenetics, ecological modelling

## Abstract

The incorporation of ecological processes into models of trait evolution is important for understanding past drivers of evolutionary change. Species interactions have long been thought to be key drivers of trait evolution. However, models for comparative data that account for interactions between species are lacking. One of the challenges is that such models are intractable and difficult to express analytically. Here we present phylogenetic models of trait evolution that includes interspecific competition amongst species. Competition is modelled as a tendency of sympatric species to evolve towards distinct niches, producing trait overdispersion and high phylogenetic signal. The model predicts elevated trait variance across species and a slowdown in evolutionary rate both across the clade and within each branch. The model also predicts a reduction in correlation between otherwise correlated traits. We used an Approximate Bayesian Computation (ABC) approach to estimate model parameters. We tested the power of the model to detect deviations from Brownian trait evolution using simulations, finding reasonable power to detect competition in sufficiently large (20+ species) trees. We applied the model to examine the evolution of bill morphology of Darwin’s finches, and found evidence that competition affects the evolution of bill length.

## Introduction

There is an increasing drive to combine evolutionary and ecological perspectives in order to fully capture the long-term dynamics of ecological communities (Johnson & Stinchcombe 2007, Cavender-Bares et al. 2009, Schoener 2011, Pennell and Harmon 2013, Hadfield et al. 2014, Price et al. 2014, Pigot and Etienne 2015). This has led to insights into the roles of ecological processes such as competitive exclusion and character displacement in shaping distributions of traits (Webb et al. 2002, Kraft et al. 2007, Emerson and Gillespie 2008, Vamosi et al. 2009). However linking such patterns in data to underlying processes is difficult, since any given pattern could be the outcome of several processes (Dayan and Simberloff 2005, Mayfield and Levine 2010).

Evidence that competition has shaped trait evolution has been generated using two main approaches. This first is the observation of character displacement, i.e. a tendency for species with overlapping ranges to exhibit increased phenotypic differences where they coexist (Schluter and McPhail 1992, Dayan and Simberloff 2005, Pfennig and Pfennig 2010, Stuart & Losos 2013). The second source of evidence for competitive effects makes use of a phylogeny to measure the distribution of species trait values relative to a null model (Webb et al. 2002, Freckleton and Harvey 2006, Vamosi et al. 2009). This is especially useful for adaptive radiations, where typically several similar species are confined to the same geographical area. Distributions that are more even than expected by chance (Webb et al. 2002, Dayan and Simberloff 2005, Davies et al. 2012) are taken as evidence that past competition caused species to seek unique ecological niches.

Convergent evolution of sets of species in separate clades has also been observed and interpreted as evidence of interspecific competition (Moen and Wiens 2009). With close niche packing interspecific competition can reduce evolutionary rates, even with a changing environment (De Mazencourt et al. 2008). Phylogenetic comparative models of adaptive radiations have slowing evolutionary rates, implicitly assuming that competition for finite niche space is an underpinning mechanism (e.g. the ‘early burst’ model, Harmon 2010a). Despite much study, however, the importance of competition remains uncertain (Gillespie 2001, Cavender-Bares et al. 2009) and, importantly, direct tests for evidence of past competition in phylogenetic data are lacking.

One approach could be to explicitly model the evolution of traits in systems of species in which competition is occurring. In general evolutionary models use some combination of continuous random change through time (Felsenstein 1973), possibly with changes of rate (Garland et al. 1992, Pagel 1997, Freckleton et al. 2002, Blomberg et al. 2003, Eastman et al. 2011, Revell et al. 2011, Thomas and Freckleton 2012), discrete random changes at speciation events (Ingram 2010), or shifts in shared adaptive optima (Uyeda and Harmon 2015). However phylogenetic models of trait evolution are ecologically neutral, since they are stochastic models that depend on the independent evolution of each species to be statistically well behaved (Pennell and Harmon 2013). Processes such as competition between species are typically not accounted for. In previous models species interactions have been assumed to generate phenomenological outcomes. For example models may assume rate slowdowns associated with competition among lineages either implicitly by modelling through time (Harmon et al. 2010) or explicitly (Mahler et al. 2010). Several models include clade-wide non-random effects (Felsenstein 1988, Hansen 1997, Price 1997, Harvey & Rambaut 2000, Freckleton & Harvey 2006, Bartoszek et al. 2012), reflecting the interaction of species with their environment, but none of these models permits trait values to be influenced by interspecific interactions.

Phylogenetic data has been simulated with competitive interactions (Freckleton et al. 2003, Nuismer and Harmon 2015). However, direct parameterisation with data is difficult because of the complexity of accounting for interspecific interactions. Niche-filling models of trait evolution on trees (Price 1997, Rambaut and Harvey 2000, Freckleton and Harvey 2006) are models of adaptive radiations where new species move discretely to the nearest of a random set of points (niches) in trait-space. Simulations under these models show that such ecological processes affect inferences drawn from comparative analyses. The most important conclusion from the analysis of such models is that methods based on Brownian motion are inappropriate, or even misleading, when applied to traits evolving in such systems. However the problem of modelling such data has never been satisfactorily resolved (Freckleton and Harvey 2006), largely because of the complexity of statistically describing the traits of a set of interacting species.

In terms of fitting complex models to data one potential approach is Approximate Bayesian Computation (ABC; see Beaumont 2010). This provides a simple method for generating posterior probabilities of models, provided we can simulate them. It is therefore well suited to fitting complex models, where it is not possible to compute a likelihood function. In this way species interactions can be incorporated into evolutionary models, thus permitting better inference of the underlying ecology. ABC has been explored for simple phylogenetic trait evolutionary models (Kutsukake and Innan 2013) including birth-death models (Slater et al. 2012), but its flexibility has not previously been used for including complex effects like interspecific interactions.

In this paper we introduce a new model for the evolution of interacting species within phylogenetic data. Our objective is to create a model that includes interactions and makes realistic predictions, but that also may be fitted to real data. The model predictions are compared with those of the Brownian motion and rate-change models for sympatric clades. We then outline how ABC methods may be used to detect competition effects and we show that that the model is readily fitted to data. Finally, we apply these methods to a simple case study, the adaptive radiation of Galapagos finches (*Geospiza spp*).

## Methods

### The model

Under the Brownian motion (BM) model of trait evolution (Felsenstein 1973), for each species *i*, a trait value *x_i_* evolves according to the differential equation,

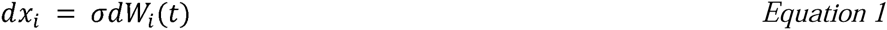
where *W*(*t*) is the integral of the continuous white noise function, such that over a finite time it has a normal probability distribution: *W*(*t*)~ *N*(0, Δ*t*). The BM model has two free parameters, the evolutionary rate σ and the root trait value. The expected variance between tips is proportional to the branch length separating them.

Most models for comparative data are based on modifying this model by adding additional parameters (Pagel 1997, Blomberg et al. 2003, Venditti et al. 2010, Eastman et al. 2011, Revell et al. 2012, Thomas and Freckleton 2012). Notably such models assume that the evolutionary trajectories of species traits are independent and do not account for interactions between different species.

Our competition model is based on the BM model, with a term added to account for interspecific interactions. Competition is modelled such that species with similar trait values tend to evolve away from each other, while species with dissimilar trait values have little influence on each other. To achieve this we assume a flat fitness surface for trait values in the absence of other species. In effect we assume that if the trait in question has a one-to-one correspondence with some resource, e.g. body size and prey size, then the distribution of resources is flat. We assume that a species with a given trait value has a corresponding ‘ideal’ resource but also uses up other resources such that the distribution of resource types used is normal and centred on the ‘ideal’ resource type. Therefore a Gaussian curve is associated with each species along a single trait axis represents this resource use and consequently its amount of influence on other species as a function of the difference in trait value between the two species (Doebeli and Dieckmann 2003, Pigolotti et al. 2010, Liemar et al. 2013, Liemar et al. 2008).

The repulsion between two species in trait space is assumed to be proportional to the overlap of each of their associated curves. For the evolution of a single trait *x* in a species *i*, we get a deterministic term, scaled by a parameter *a*, in addition to the BM term:

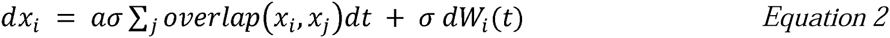

The area of overlap of two normal curves is related to the cumulative normal function *Φ* of minus the distance (in standard deviations) between them, such that the overlap is equal to 2 *Φ(−distance* / 2) (Inman and Bradley 1989). The overlap of two curves very far from each other is 2*Φ*(−∞) = 0, whereas the overlap of two curves with the same centre is.

The relative intensity of competition is measured by the competition parameter *σ* and the Brownian rate parameter in combination. Ideally we would have chosen to make the kernel width an additional parameter of the model. However in practical terms it would not have been possible to distinguish this effect from that of the competition parameter *a*. Appendix A shows that to a linear approximation the effects of the two are the same, and so they are likely to be statistically indistinguishable.

The instantaneous change of the trait value *x_i_* of species *i* is given by equation 3:

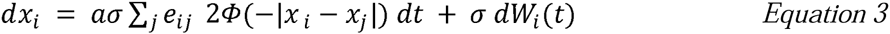

Each*x_j_* is a vector in trait space; the index *j* denotes species. The right-hand-side has two terms: the first is a deterministic ‘competition’ term, which pushes apart species that are nearby in trait-space. *e_ij_* is the unit vector pointing from species *j* to species *i* in trait-space. Thus, *a e_ij_ Φ*(−|*x_i_* − *x_j_*|) is a vector in trait-space pointing from species *i* to species *j*, proportional to the model parameter *a* and depending on the closeness in trait-space of species *i* and *j*. What distinguishes this model from previous ones is that in the competition term of the equation all traits are linked: the evolution of two species away from each other in trait space depends on the Euclidian distance between them, as well as their distances to all other species.

We largely concentrate here on single resources and traits. However, more generally a multivariate normal curve in trait-space may be associated with each species in order to model interactions along several resource axes.

In both the BM and competition models, trait variance increases without bound as time progresses. In reality there are limits that will be driven by ecology or by developmental and physiological constraints. We therefore adapted the model by imposing hard limits on trait-space, such that species can evolve up to a chosen extreme value but no further. This model was simulated alongside the limitless model and hence we obtain a new model with constrained trait/niche space. We assume that the limits are symmetric and defined by the most extreme value *L*.

### Simulation framework

In a BM model trait evolution may be modelled readily and quickly because species are assumed to be independent. However our model requires that we simulate evolution over interacting branches, which adds to the computational overheads. The approach used here is to take slices of evolutionary time separating speciation events and to apply our model of evolution to each slice sequentially. Within each slice branches are evolved simultaneously. Simulation starts at the root of the tree, with time (branch length) *t* to the first speciation event. At speciation the trait value of the speciating branch is copied to both of its offspring. The time to the next speciation event is found, and the process is repeated until we have the required tip trait values, including a period between the final speciation event and the tips.

Simulations were performed on random ultrametric trees generated under a Yule process (TESS, Hoehna 2013) with 20 to 80 tips. These tree sizes are typical for datasets used to test for evidence of adaptive radiation. Because the competition model is designed for sympatric, interacting sets of species having undergone adaptive radiation, it is unlikely that numbers of species will be very large. Values of trait variance across tips were calculated as the mean of the tip variances of 1000 trees with the same tip count.

The same routine was used to compute the correlation between correlated traits under different competitive regimes, this time with a fixed tree size of 50 tips. We also generated distributions of tip trait values, for Brownian models without competition, and with and without limits. For a single random phylogeny, each evolutionary model was simulated 1000 times, and all the tip values pooled to create a frequency distribution. We repeated the same simulations for trees of different sizes (20, 40, 60 and 80 tips) and calculated the variance of each set of tip trait values. This enables the assessment of how the evolutionary model affects the relationship between phylogeny size and trait variance.

### Model comparisons and likelihoods

We fitted the model to data using Approximate Bayesian Computation (ABC) (reviewed in Beaumont 2010, Csilléry 2010, Hartig 2011). ABC can be used for comparing the probabilities of datasets under different models when these probabilities are difficult to compute directly. This is because the only requirement to perform ABC is that we can simulate new datasets using the model. The dataset probabilities are approximated by simulating a large number of datasets, and ‘accepting’ only those simulations that are very similar to the observed dataset. This similarity can be judged either from the data values themselves, or using summary statistics. The proportion of simulations that are accepted is then assumed to be proportional to the dataset probability. When the model contains continuous parameters, we sample across these parameters and obtain an approximate probability density for the observed data under any point in a range of parameter values. This can be used to estimate the likelihood curves of fitted models. ABC relies on the likelihood being a fairly smooth function of the model parameters (Hartig 2011).

To apply ABC to phylogenies (e.g. Beaumont 2010) we sample the parameters of the evolutionary model randomly many times from a prior parameter distribution. Here we choose the prior distribution to be uniform, with the model necessitating a hard limit at zero for both the Brownian rate and the competition strength. For each set of parameters, trait datasets are then simulated for the known phylogeny. Summary statistics are generated for the simulated data, and only those simulations for which the summary statistics are within a small value *ε* of the observed data’s summary statistics are accepted. Thus, for observed data *D* and tolerance*ε*, we accept some simulated data *D′* if

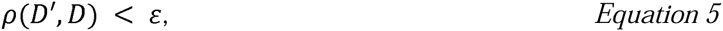
where *ρ* is the discrepancy, or distance in summary statistic space, between *D*′ and *D*. In practice we chose the tolerance *ε* based on the size of posterior sample that we wanted to obtain, so we might simulate a million datasets and choose *ε* such that we accept the best 500 simulations. By plotting acceptance rate against parameter values, we get an estimated likelihood surface.

To compare simulated and observed datasets, we need to compare summary statistics. We chose to use three summary statistics: the mean and the variance of the differences between each species and its closest neighbour in trait space, and the overall phylogenetic signal as measured by Blomberg’s *K* (Blomberg et al. 2003). The rationale for using these three was to capture the overall amount of evolution, the overdispersion of trait values, and the phylogenetic structuring of the trait values. There is no well-established procedure for choosing summary statistics for ABC. High sufficiency is needed to compare models, but the ABC method quickly loses accuracy and stability with large numbers of summary statistics (Csilléry 2010). Our summary statistics were chosen on a pragmatic basis, since they capture the important aspects of the model’s behaviour, namely increased divergence between sibling species, and an even overall distribution of traits across the phylogeny. Hartig et al. (2011) recommend empirical tests of sufficiency by comparing analyses that reject using summary statistics to ones that use the actual data. We performed this test for small trees (5 tips) with simulated datasets, and found that the three summary statistics resulted in power and parameter estimates that were not distinguishable from those produced by using the tip trait values themselves as the summary statistics used to compare models and generate likelihood estimates.

We chose to compare the competition model with the BM model using maximum approximated likelihood, because the BM model is embedded in the competition model. The null and alternative ABC acceptance rates *A* give an estimate of the likelihood *L(H/D)* of the observed dataset under the various model parameters, assuming a smooth probability distribution. Then the log-likelihood ratio statistic for the comparison of two models *H*_0_ and *H*_1_, when there is no prior difference in model likelihood expectation, is given by:

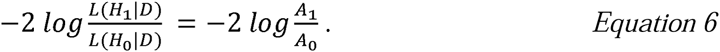

When the models are nested, this test statistic approximates *χ*-squared distribution given certain assumptions: large samples and normally distributed parameters. However, these assumptions may be significantly violated by phylogenetic methods (Freckleton 2009). For instance, in the BM model *σ* is bounded at zero, and in our competition model *a* is also bounded at zero. To correctly interpret the test statistic, therefore, the null distribution of the log-likelihood ratio test statistic was assessed with a parametric bootstrap.

The bootstrap was undertaken by performing the model comparison analysis on datasets generated under BM, to create a null distribution of likelihood ratios. Then if, for example, we want to know the likelihood ratio corresponding to a p-value of 0.05, we simply look at the 95^th^ percentile of the null distribution. The resulting Type I error rate is therefore chosen by design: if a significant likelihood ratio is one that corresponds to a p-value of 5%, then the Type I error rate is 5%. To estimate typical significance thresholds, we performed this procedure for random trees using 1000 random datasets.

The power to reject BM in favour of the competition model was assessed by using random ultrametric Yule trees (20, 40, 60 and 80 tips). The bootstrap process was performed to determine the significance threshold for that tree. Then, for a given value of the competition parameter *a*, we simulated a large number of datasets and determined the likelihood ratio (between the BM and competition models) for each one. The proportion of these datasets which showed significant support for competition effects defines the power of the model for that value of *a*. We repeated this process for a range of competition strengths from to *a* = 0 to *a* = 5. This range covers evolution from a Brownian process with no interspecific interaction (*a* = 0) to a largely deterministic regime with high phylogenetic structuring of trait values *(a =* 5).

To evaluate the simulated data produced by the competition model, other comparative models were fitted to the data: the Brownian model itself, the *κ*-model, which measures the degree to which evolution is speciational rather than gradual (Pagel 1997), and K, a measure of phylogenetic signal (Blomberg et al. 2003). Parameter estimates were generated using the R packages geiger (Harmon et al. 2008) and picante (Kembel et al. 2010).

The dataset simulations were written in C++. Scripts for using these datasets for likelihood estimation were written in R (R Development Core, 2005), using ape (Paradis et al. 2004) and TESS (Hoehna 2013) for tree generation (code available online).

## Results

### Example of clade evolution under the competition model

Illustrative examples of evolution under the competition model are shown in Figure 1. Estimates of phylogenetic signal and of rate-change transformations were generated for these simulated data and also are shown in Figure 3. The evolution of each species is tracked through time from left to right.

**Figure 1.**
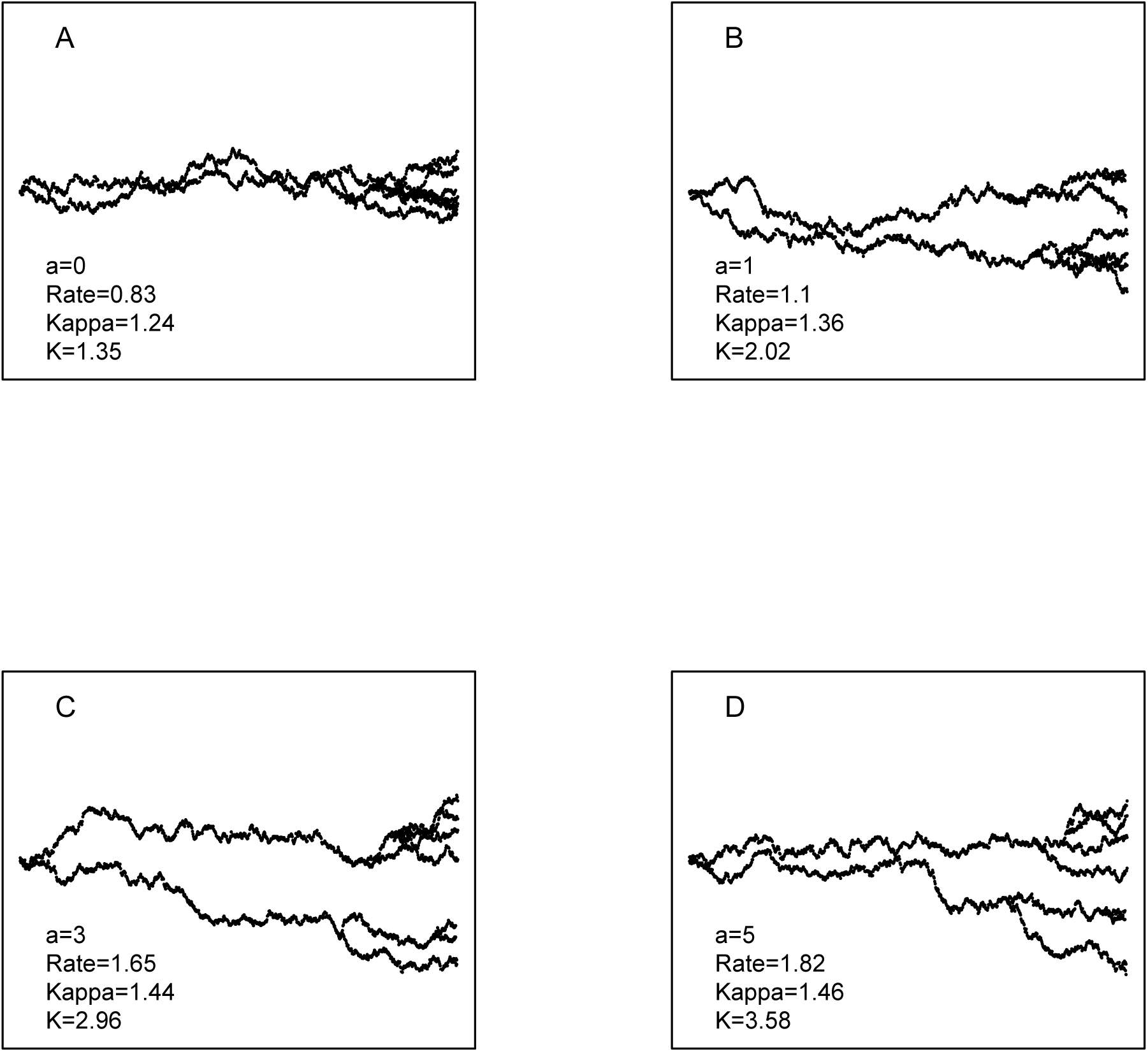
Examples of a single trait evolving under the competition model with different strengths of the competition effect parameter *a*. The parameter values listed by each plot are the averaged estimates under those respective models for several datasets generated under the competition model. When *a* = 0 we recover the nested BM model.

Competition tends to increase the overall variance in traits amongst the species in a phylogeny, as is clear from the increase in range and variation of traits moving from Figure 1A to Figure 1D. This is because species experiencing competition from other species are more likely to evolve extreme trait values to become more different and escape competition.

As the strength of competition is increased, the differences between species become more clearly defined, with them occupying distinct positions in niche space. There are fewer crossings over of trait’s evolutionary paths over time between species, and the phylogenetic signal *K* exceeds the neutral BM prediction of *K =* 1. Competition thus increases phylogenetic signal above that expected under the BM model, while presenting the appearance of a considerable tree-wide evolutionary slowdown. This means that a species’ trait values map more directly onto its position in the tree. For sympatric clades, there is thus a prediction of traits being more phylogenetically conserved than under BM.

Typical estimates for commonly used branch transformation parameter *κ* from these datasets are also shown in Figure 1. *κ* measures the rate change along branches, and overall measures the degree to which change is speciational (Pagel 1997). A transformation parameter *δ* models the overall changes in evolutionary rate across the tree, with lower values corresponding to evolutionary slowdowns (Pagel 1997). We find that the *δ* parameter diminishes very rapidly as competition is increased. This reflects an apparent slowdown of evolutionary rate, which becomes more pronounced as the value of *a* increases. Species competing for unoccupied niche-space thus evolve more rapidly early on in their development, when they are more similar to one another and the effects of competition are stronger, as one would expect in an adaptive radiation (Yoder 2010).

### Trait distributions across tree tips

The distribution of trait values of the phylogeny tips is flattened in the competition model compared with BM models, which predict normal distributions for large trees. This outcome is expected when competition shapes trait values (Davies et al. 2012). Figure 2A shows the model predictions for tip trait value distributions. The impact of competition on trait distributions is even more pronounced where hard limits are placed on the available range of trait values.

**Figure 2.**
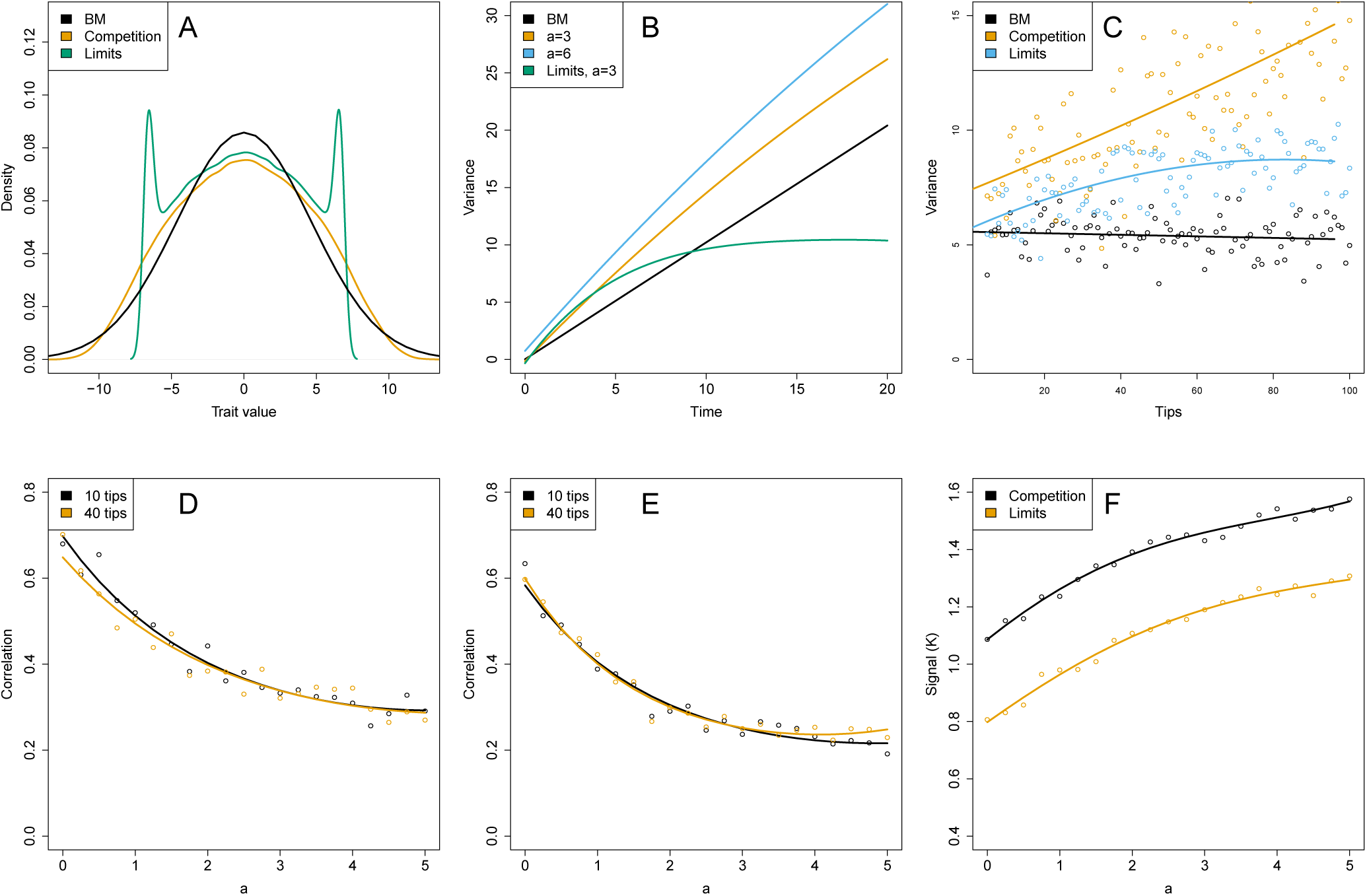
A, B, Effects of interspecific competition on trait value distributions across a clade. The BM model results in a normal distribution, whose variance increases linearly with time. The competition model predicts a flattened distribution, whose variance increases with time, initially faster than for BM, but slowly slowing to the BM rate. When hard limits are placed on trait space, the competitive distribution is further flattened, with probability peaks at the limits, since competition will tend to push species to those limits. C, Effect of tree size on variance of trait values of an evolving clade. Under BM the tree size has no effect, but with competition across the clade, a more numerous clade results in a greater amount of trait variation in that clade. D, The correlation between two traits across the species in a clade, as a function of the strength of interspecific competition parameter. The BM rate parameter in set to 1 throughout. E, Correlation for the model with limits. F, The signal (Blomberg’s *K*) as a function of competition strength. The traits are evolving with their random (BM) evolution strongly correlated; the pressure from competition to be dissimilar acts against this natural correlation.

In addition to creating a more even trait distribution, competition increases the overall amount of trait divergence, given equal BM rates (Figure 2B). This is consistent with the expectation that equivalent species sets should be more diverged in sympatry than in allopatry when there is competition (Schluter 2000). From a biological perspective there is thus a prediction that competition leads to a wider range of morphological variation in a clade, reflecting the increased tendency towards extreme traits when there is lots of competition.

### Effects of tree size

We used trees normalised to the same total length, regardless of the number of tips. Under BM and rate-change models, the variance of tip trait values shows no change with increasing the number of tips (in agreement with Ricklefs 2004). In the competition model larger trees have greater variance, since a greater number of species are ‘pushing’ each other away; this is shown in Figure 2C. This relationship seems to be approximately linear for the unbounded competition model. When hard limits are imposed, the variance reaches a maximum corresponding to the position of the extremes.

### Effects of competition on correlated traits and phylogenetic signal

For pairs of traits, in which the evolutionary changes in trait values are correlated, the correlation between the traits decays rapidly with increasing competition strength. This is even more pronounced when there are limits on extreme trait values. Figure 2D and Figure 2E show how the correlation decays. By de-correlating traits, competition forces the trait space to be occupied more evenly. An interesting corollary to this may be an observable pattern of high correlation between traits within species, but reduced correlations between species.

Phylogenetic signal is increased by competition as species tend to be remain adjacent in trait space to their close relatives (Figure 2E), since their trait values are unlikely to ‘cross over’ with time. Plots of traits through time therefore become more defined and tree-like. This can be seen for example in the sample simulations of Figure 3. Correlation between traits has little effect on the phylogenetic signal exhibited by the individual traits under either the BM model or the competition model. Limits reduce the phylogenetic signal, since there is less trait-space for distantly related species to diverge. Indeed, without competition driving the signal up (i.e. when *a =* 0), the model with limits predicts reduced signal compared with the BM model, with *K* < 1.

**Figure 3.**
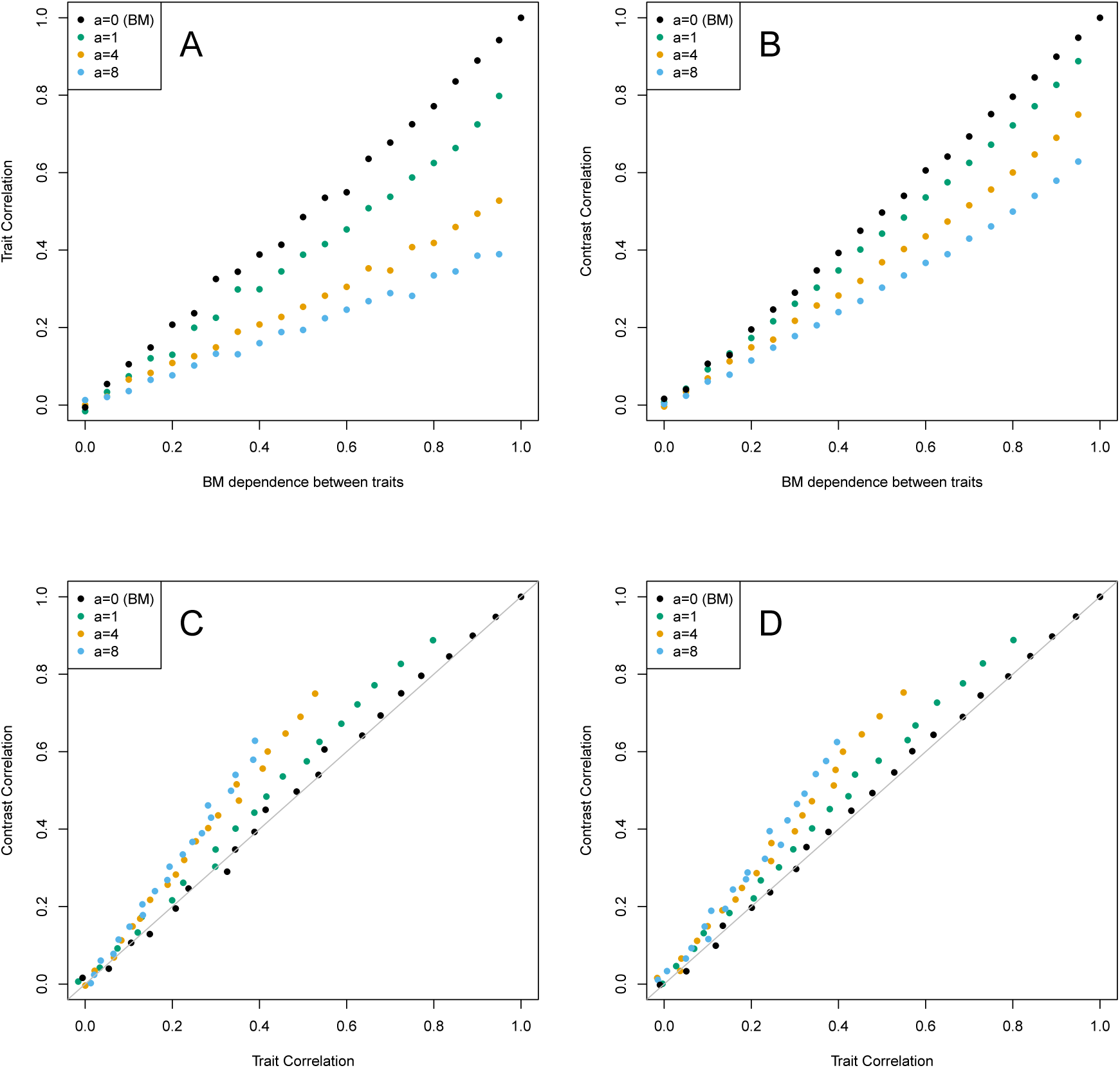
Traits and contrasts for two traits, where one trait has a dependence on the other. For each step in time, the dependent trait has an evolutionary change that depends on the change to the other trait. If the dependence is 1, then these changes are equal; if the dependence is 0.5, then the change in the dependent trait is 0.5 of the change in the other trait, and the remaining change is random. A, Trait correlation as a function of the intrinsic trait dependence. B, Contrast correlation as a function of the intrinsic trait dependence. C, Correlation between contrasts is slightly greater than correlation between traits for competitive evolution. D, Contrast and trait correlations for a model of competition with traitspace limits.

Price’s (1997) model of adaptive radiations has the feature that when two traits have correlated evolution, the phylogenetically independent contrasts are less strongly correlated than the traits themselves (Price 1997, Rambaut and Harvey 2000, Freckleton and Harvey 2006). A Brownian model predicts equal correlation for both traits and contrasts.

We compared trait and contrasts correlations under the competition model presented here. Competition tends to reduce correlation between traits, as discussed above, but we set the Brownian evolution of the traits to have very high correlation. The results are shown in Figure 3. In contrast with the result for the Price (1997) model, we found that contrasts had slightly higher correlations than traits. This probably reflects the fact that competition tends to have a greater effect earlier in the evolutionary history of any particular species.

### Power

The power to detect competition effects against a background of BM evolution is shown in Table 1 for trees of various sizes. We define the power as the frequency with which simulated datasets show significant support for competition effects as opposed to the (nested) BM null model. Power is greatest for large trees with high competition strength relative to BM rate. This can be interpreted as the relative contribution to overall evolutionary change of competitive effects versus other, effectively random, effects.

**Table 1.**
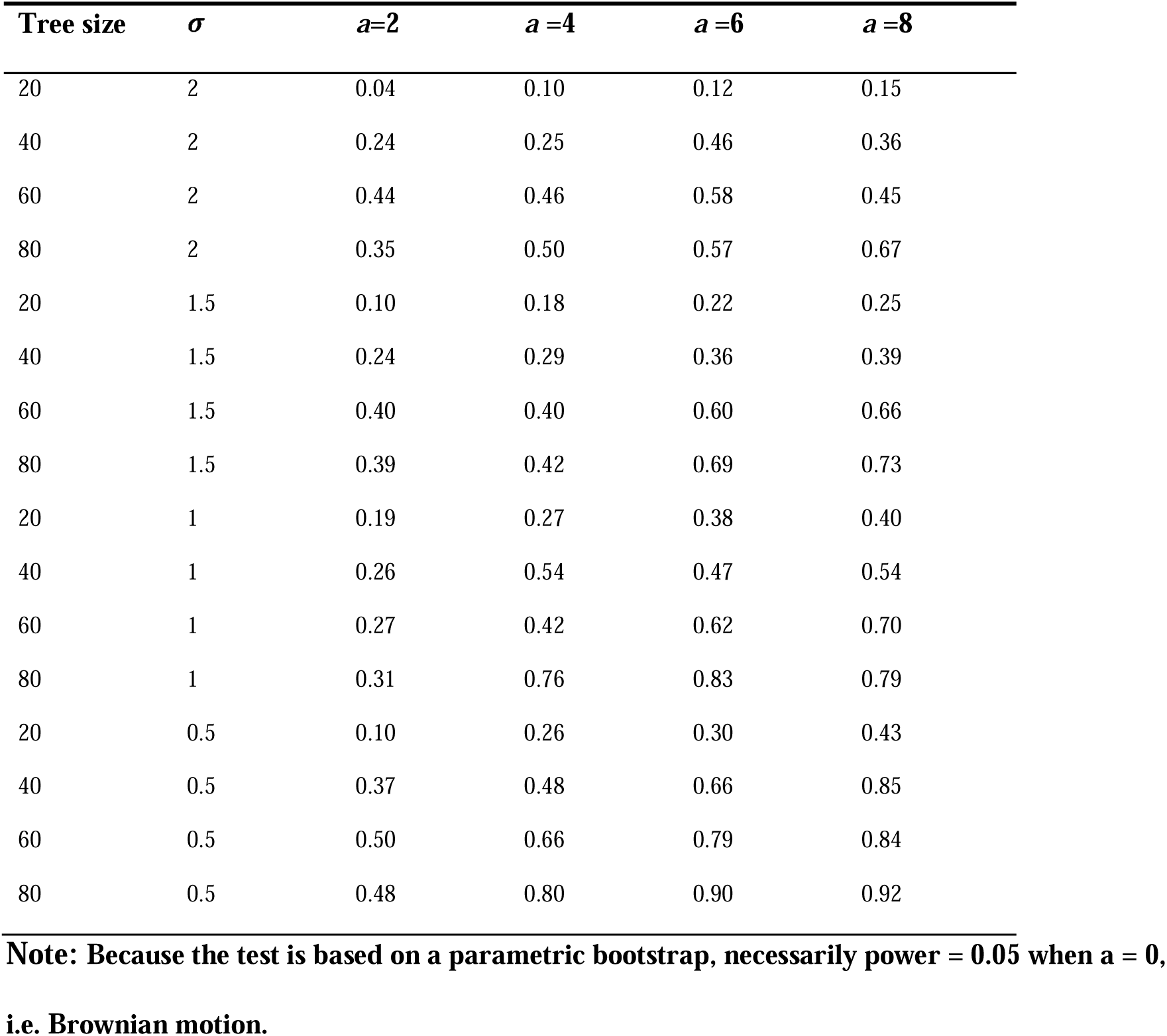
Power to detect competition versus Brownian Motion as tree size, the Brownian parameter *(σ)* and intensity of competition (*a*) are varied.

In this context a significant dataset is one for which the Type I error rate is estimated to be ≤ 0.05. This is the frequency with which data from null model simulations display model likelihood ratios that equal or exceed the ratio for the ‘observed’ dataset. This is determined via a parametric bootstrap.

## 4 Case study: Galapagos finches

The simulations described above demonstrate two things: first that the model we describe successfully captures behaviour that we would expect to be observed in systems of interacting species. And second, that it may be applied to data and used to infer the presence of competitive interactions. In order to use the model in a real-world example we applied the competition model to an example dataset, using trait measurements collected in Harmon et al. (2010a; originally Grant and Grant 2002, Lack 1947; repository in Harmon 2010b http://dx.doi.org/10.5061/dryad.f660p), and a recent molecular phylogeny (Lamichhaney et al. 2015). We used the Galapagos finches *(Geospiza spp.)*, since they are a well-studied adaptive radiation, and ecological effects were anticipated to be of importance. The effect of character displacement on intraspecific variation among these finches is well documented (e.g. Grant and Grant 2006). Here we are looking to see whether the same mechanism has had an effect on the overall distribution of traits across the clade.

### Methods

The phylogeny of Galapagos finches was taken from Lamichhaney et al. (2015). Data was available for five traits: wing length, tarsus length, bill length (culmen), bill depth and bill width (gonys) (Harmon 2010b). We computed likelihood ratios for each trait individually.

After simulating data on the phylogeny to determine likelihood cutoffs for rejecting BM, a likelihood comparison between the competition model and the nested BM model was run for each of the five traits separately. We performed the tests twice, once including and once excluding the phylogenetic summary statistic *K*, to judge the importance of signal in favouring the competition model. Applying the bootstrap process for single traits gave a likelihood ratio for the full tree of 3.49 corresponding to p=0.05, while 3.79 corresponded to p=0.01. There is one degree of freedom difference between the competition and BM models.

### Results

The parameter estimates and model likelihood ratios are shown in Table 2. The beak shape traits showed greater support for competition compared with Brownian evolution than the other traits. This appears to point to an ecological effect: the competition model implies a tendency towards well-differentiated niches that don’t cross, and the beak shape is an ‘ecological’ trait, in the sense that it corresponds strongly to feeding habits (Grant and Grant 2011). Figure 4 shows illustrative plots of simulated trait evolution using the model parameters that were estimated for the culmen length. Compared with BM, shown in Figure 4A, the tree becomes very well defined, with strong phylogenetic signal.

**Table 2:**
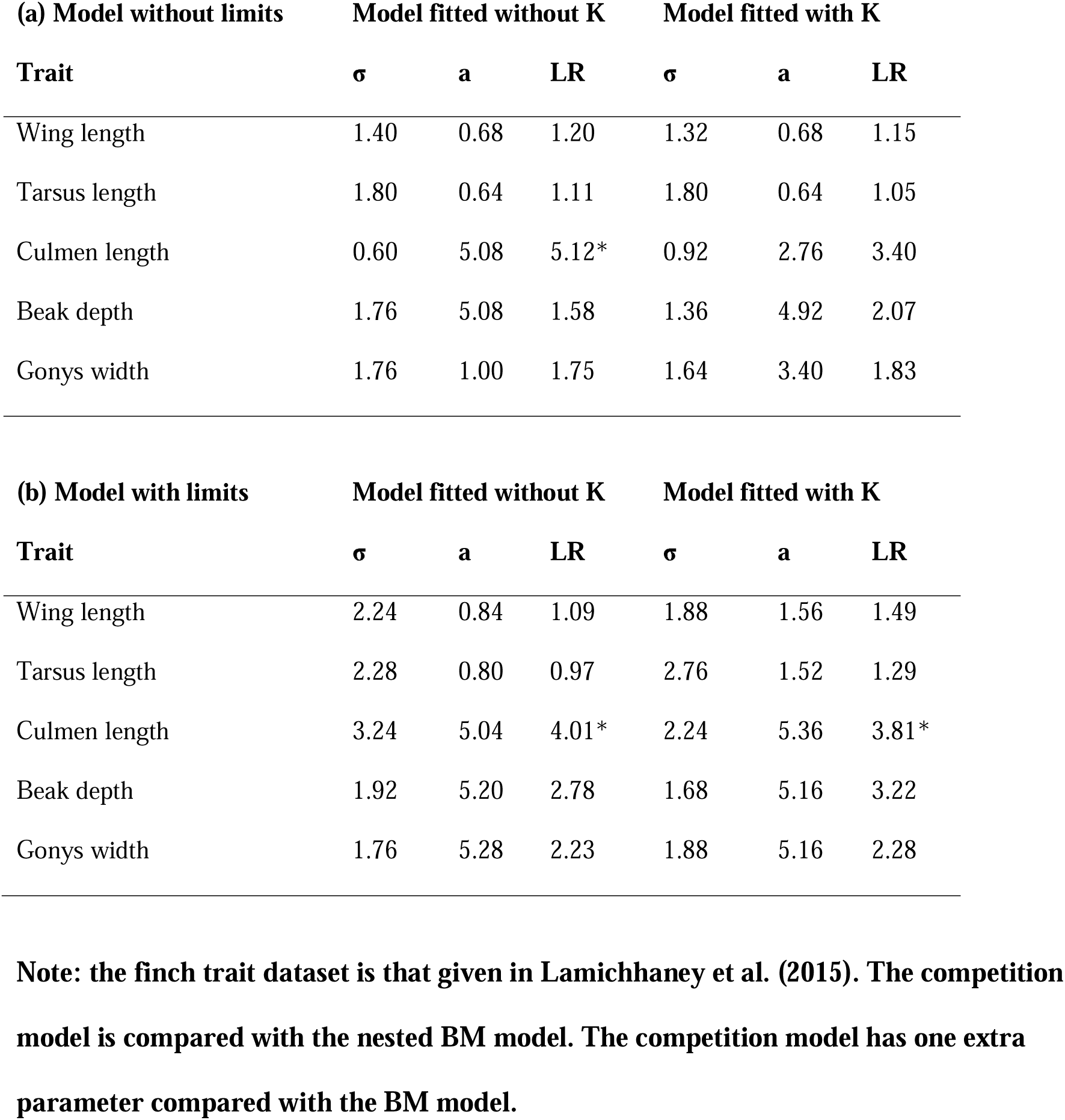
Traits and likelihood ratios for model comparisons for Galapagos finches.

**Figure 4.**
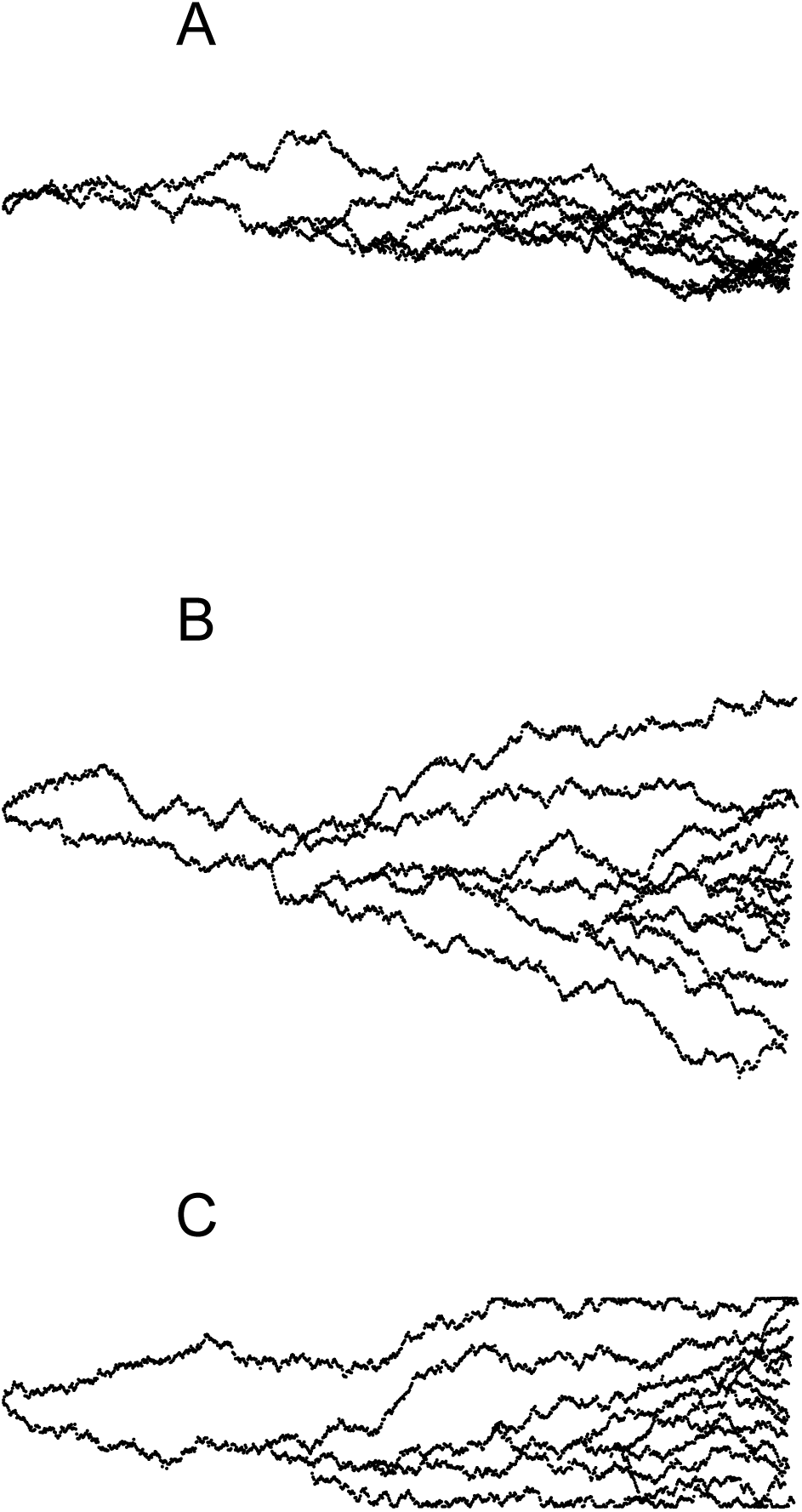
Example simulations of univariate trait evolution on the Darwins finch phylogeny, using model parameters estimated from culmen length data. A, Evolution under Brownian motion. B, Evolution including the competition parameter. C, Evolution including competition and hard limits

One of the beak traits, culmen length, favoured the competition model when signal was not used, but less so when it was included. Brownian rate parameter estimates under the model with limits are higher than those for the non-limited competition model. This higher rate does not result in a greater total amount of evolutionary change, due to the hard limits that are reached either way. This result does, however, suggest that the niche landscape may be the limiting factor in the finches’ evolution: the finch traits are capable of evolving rapidly, but their values are constrained by the combination of interactions between species and environmental limits on niche space.

It is worth noting that none of these results takes into account intraspecific variation or uncertainty in the finch phylogeny. Species interactions will be somewhat independent on different islands, as illustrated by the character displacement seen in intraspecific trait variation (Grant and Grant 2006). Furthermore, the tree topology is fairly uncertain, since many of the tree node dates have error bounds that are large compared with the distances between nodes (Lamichhaney et al. 2015).

## Discussion

There have been several recent approaches to integrating ecological processes into phylogenetic models of evolution (Cavender-Bares et al. 2009, Hadfield et al. 2013, Pennell and Harmon 2013). As a step in this direction, we have created a model of interspecific competition on phylogenies of coexisting species. The model provides a process-based picture of competitive evolution, linking statistical patterns directly to the underlying ecology. It generates the patterns we expect to see in situations where interspecific competition is important.

Competition and niche overlap have a complex relationship, and the former may cause or inhibit the latter depending on the particular niche being measured. According to one possibility, there is an optimum position on a niche axis, and species will compete with each other to occupy it. Consequently they will evolve similar trait values tailored to that optimum (Colwell and Futuyma 1971). Another possibility is that the niche is represented by a continuum of equal fitness along the axis. In this case species will compete for empty regions of the niche axis and evolve minimally-overlapping, evenly-spaced trait values, consistent with the ecological idea of character displacement (Grant 1972, Strong et al. 1978, Dayan and Simberloff 2005). Our competition model accommodates the latter in a phylogenetic context.

One prediction of the competition model is a flattened distribution of trait values among contemporary species of a single sympatric clade, in common with several previous models. Indeed, competition is often inferred from such evenness (Dayan and Simberloff 2005, Davies et al. 2012). The same pattern can however be caused by competition at the community assembly level rather than in situ trait evolution (Cavender-Bares et al. 2009, Stuart and Losos 2013), or by geographical structure in speciation and extinction (Pigot and Etienne 2015). For this reason the model presented here is applicable only to clades known to share a common environment, or subsets of clades formed by restricting attention to a single environment. Since the model incorporates pairwise interactions between all the species present, it requires that each species be at least somewhat sympatric with each other species. Strong niche-conservatism is then predicted for sympatric clades. However, it is worth commenting that species within a clade are not exclusively sympatric or allopatric in their distributions. Across clades there is a wealth of variation in species’ geographic structure and opportunity to interact (Fitzpatrick et al. 2008). Taking variation in range overlaps into account may be an interesting future development in such modelling.

The pattern of non-Brownian value distributions and high phylogenetic signal is also generated by an alternative, but related, mechanism, where instead of a continuously available space of niches, niches are discrete, with one species per niche, and new species arise in nearby discrete niches (Price 1997, 2014). Determining a method to distinguish this model will be a key advance. Specifically it differs from the current model in that the tree topology is a function of trait values and not fixed as in our model.

Most phylogenetic models of trait evolution are modifications of the random BM model. As noted above, adaptive radiations are generally consistent with a tree-wide gradual slowdown in evolutionary rate (delta-model: Pagel 1997; ACDC model: Blomberg et al. 2003). Speciational evolution can be modelled as a gradual branch-wise slowdown (*κ* model, Pagel 1997), or by partitioning evolution into gradual and speciational parts (Bokma 2008, Ingram 2011). Discrete shifts in evolutionary rate can be modelled to detect, for example, adaptive radiations embedded in a larger tree (O’Meara et al. 2006, Thomas et al. 2006). Slowdowns in evolutionary rate have also been observed as a function not of time but directly of a clade’s size (Mahler et al. 2010). The results for our competition model suggest that it reproduces the appearance of a strong tree-wide slowdown. During a radiation, though, competition is predicted to cause overall trait variance to increase much more rapidly. Our results for the competition model also demonstrate raised phylogenetic signal, in agreement with similar results in Nuismer and Harmon (2015).

In all analyses we used a fixed competition kernel width. The fact that this width is not distinguishable from the competition strength itself suggests that the amount of variation possible within a single niche is not readily ascertained from a phylogeny and trait data. Measurements of intraspecific variation will be more suited to this question. In fact, the competition kernel widths could be set empirically before analysis, if data on intraspecific variation were available.

Our results for the Galapagos finches support the well-known presence of character displacement in that clade (Grant and Grant 2006), and further suggest that interspecific competition is a significant force comparable with other, effectively random, sources of evolutionary change for the Galapagos finches. For some beak shape traits, the Galapagos finches exhibit the elevated phylogenetic signal predicted by the competition model.

The usefulness of the model in detecting competition in other datasets will depend on the strength of competition effects in nature. The model could readily accommodate known extinct species, but unknown extinct species may affect the contemporary datasets generated by the model, since we will not be accounting for their past interactions with extant species. Additionally, because it involves between-species interactions, the model is particularly sensitive to missing data, both species themselves and their trait values.

There are numerous speciation/extinction models for phylogenies (Nee et al. 1994, Pybus & Harvey 2000, Rabosky 2006, Freckleton et al. 2008, FitzJohn 2010), including some that are expected to correspond to clades with interspecific competition (Harmon et al 2010, Etienne et al. 2012). Our model is concerned only with trait evolution. Trait evolution and diversification rates may be coupled in nature, however, and may both vary with factors such as interspecific competition. Building models of adaptive radiations that simultaneously predict trait evolution and diversification will be key in the future.

As phylogenetic methods continue to be used to infer evolutionary processes, it will be important to include specific ecological mechanisms (Vamosi et al. 2009). Competition for ecologically distinct roles is often implicitly or explicitly assumed in adaptive radiations, but its prevalence and importance remain uncertain (Schluter 2000, Stuart and Losos 2013). We have developed an explicit model of competition on phylogenies, to detect competitive effects in sympatric adaptive radiations, and to enable measurement of competition strength. The predictions of this model may help to understand the roles ecological processes play in shaping trait evolution.

## Appendix: Estimating competition strength and effect width simultaneously

The overlap between species *i* and *j* is proportional to 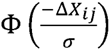where is the cumulative normal distribution. Integrating by parts yields the following approximate function for the overlap between two species:

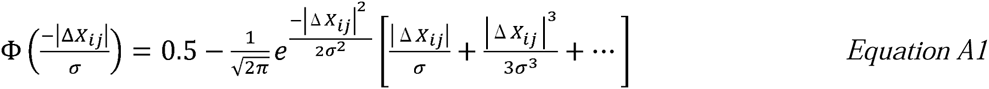

To get the evolutionary rate we multiply this by a, giving:

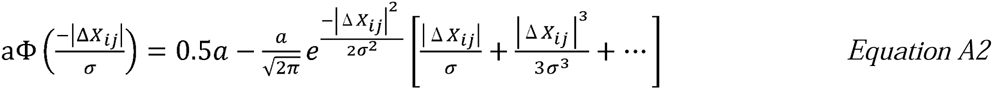

At first glance it might appear that changing *a* and *σ* would have different effects because the former changes evolutionary rates in a linear manner, whilst the effect of the latter is non-linear. However, if there are a large number of species within a limited niche space, then distances between species will be low, i.e. |Δ*X_ij_* | is small. Consequently, we can use the following approximation by the Maclaurin series expansion of *e^x^:*

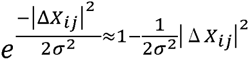

Substituting into equation (2) and ignoring higher than squared terms we get:

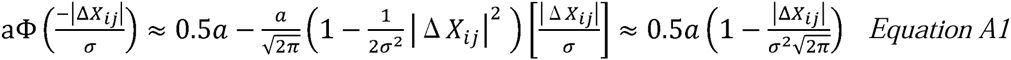

Overall, the rate of evolution is given by the overlap, 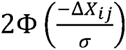multiplied by a, yielding:

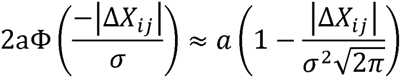

This equation is linear in both *a* and *σ*^−2^. Thus from a statistical perspective *a* and *σ* will be non-identifiable if the species are interacting strongly. If species are not interacting strongly, i.e. Δ*X_ij_* is large, then the data will contain no information on interactions between species and hence it will not be possible to fit the model and we cannot estimate either *a* or *σ*.

